# IL-33 induced gene expression in activated Th2 effector cells is dependent on IL-1RL1 haplotype and disease status

**DOI:** 10.1101/2022.10.20.513024

**Authors:** AK Saikumar Jayalatha, ME Ketelaar, L Hesse, YE Badi, N Zounemat-Kermani, S Brouwer, FN Dijk, M van den Berge, V Guryev, I Sayers, JM Vonk, IM Adcock, GH Koppelman, MC Nawijn

## Abstract

**Background:** IL-33 can activate Th2 cells after binding to the IL-1RL1/IL-1RAcP receptor. However, it is unknown whether disease or genetic make-up regulate IL33 responsiveness of Th2 cells.

**Objective:** We investigated whether *IL-1RL1* asthma risk haplotypes or asthma status influence IL-33-induced Th2 transcriptomic responses *in vitro*.

**Methods:** CD4^+^ T cells from 16 asthma patients and matched controls, stratified for *IL-1RL1* haplotype, were differentiated into Th2 effector cells, and re-stimulated in the presence or absence of IL-33, followed by RNA-sequencing and differential gene expression analysis. Association of IL-33-induced gene signatures with clinical parameters was tested in U-BIOPRED sputum transcriptomic data.

**Results:** Presence of IL-33 during Th2 cell re-stimulation results in extensive changes in gene expression, affecting a large proportion of the activated Th2 cell transcriptome. IL-33 induced more genes in Th2 cells from donors with asthma than in controls. However, the IL-33 effects were strongest in Th2 cells carrying the *IL-1RL1* risk haplotype, as evidenced by a significantly increased effect size compared to controls. A gene signature of IL-33 induced genes in Th2 cells showed a strong positive correlation with macrophage counts and pauci-granulocytic asthma in sputum transcriptomic data from the U-BIOPRED study.

**Conclusion:** The risk *IL-1RL1* haplotype strongly increases the sensitivity of Th2 effector cell for IL-33-mediated regulation of gene expression, while asthma status has an independent effect. The IL-33 gene signature was not associated with type-2 high or eosinophilic asthma in sputum transcriptomic data from U-BIOPRED.

**Clinical Outcome:** Cellular sensitivity to IL-33 depends on genetic makeup and asthma disease status.

**Capsule summary:** IL-33 strongly alters gene expression of activated Th2 cells, dependent on asthma disease status and *IL-1RL1* risk haplotype. IL-33 driven gene signature in sputum transcriptomic data identifies pauci-granulocytic asthma.

## Introduction

Asthma is a common and heterogeneous respiratory disease caused by interactions between genetic and environmental factors^1^. Asthma heterogeneity is reflected by different types of granulocytic inflammation in sputum, such as eosinophilic, neutrophilic or pauci-granulocytic^2^. Asthma genes *IL-33* and *IL-1RL1* have been associated with childhood onset asthma and type-2 high asthma^3–6^.

In response to environmental triggers such as allergen exposure, viral or bacterial infection, bronchial epithelial cells release IL-33. IL-33 acts as an alarmin, activating structural or immune cells by binding to the IL-1RL1/IL-1RAcP receptor complex. IL-1RL1 has two main isoforms, a transmembrane protein (IL-1RL1b) that is part of the heterodimeric IL-33 receptor, and a soluble protein (IL-1RL1a) acting as a decoy IL-33 receptor. *IL-1RL1* is expressed by several structural cells in the airway wall, as well as a range of immune cells involved in asthma pathogenesis including mast cells, macrophages, eosinophils, and Th2 cells^4,7^. Thus, IL-33 and IL-1RL1 can activate an immune cascade that contributes to hyperresponsiveness, remodelling and chronic type 2 inflammation of the airways^5,8,9^.

Human Th2 cells constitutively produce IL-1RL1a and express low levels of IL-1RL1b on their cell surface^10^. Th2 cells are key players in the initiation and maintenance of chronic airway inflammation in asthma^3,8^. IL-33 has been shown to promote Th2 cell differentiation of naive human Th cells and to enhance the production of Th2-cytokines such as IL-4, IL-5, and IL-13^11^. Genetic variation at the *IL-1RL1* locus affects both gene and protein expression levels due as well as signalling capacity of the transmembrane IL-1RL1 receptor, due to non-synonymous SNPs present in the intracellular TIR domain^4^. However, it is not known whether genetic variation at the *IL-1RL1* locus affects the Th2 response to IL-33. Such insights might aid the selection of patient groups that would benefit most from interventions directed at the IL-33/IL-1RL1 pathway, which have recently been shown to be efficacious in moderate to severe asthma^12,13^. Therefore, we aimed to study the functional consequences of the asthma-predisposing *IL-1RL1* haplotypes for IL-33 induced responses in Th2 cells. We hypothesize that asthma-associated *IL-1RL1* haplotypes enhance the response of Th2 cells to IL-33. Moreover, we hypothesize that transcriptomic signatures of IL-33 stimulated Th2 cells would enable selection of a subgroup of asthma patients in whom the IL-33/IL-1RL1 pathway is important.

To test our hypothesis, we performed a translational study investigating the effect of IL-33 on gene expression of Th2 cells from both asthma patients and matched controls, stratified for *IL-1RL1* haplotypes. Moreover, we tested in U-BIOPRED sputum transcriptomic data whether a signature of IL-33 induced genes could identify a subgroup of patients with type-2 high asthma.

## Methods

Detailed methods are provided in the Online Supplement.

### Patient selection, CD4^+^ T cell isolation, Th2 cell differentiation

Sixteen asthma patients were selected from the ‘Roorda’ cohort^14,15^ (METc 2012/173) and sixteen healthy subjects were selected from the ‘NORM study’^16,17^ (METc2009/007), that required no history of lung disease and normal lung function, were selected ^14,17,18^. Both groups were genotyped with Illumina Omni Express Chip^15,17^. We selected 4 asthma-associated SNPs in the *IL-1RL1* gene: one strong eQTL and pQTL for *IL-1RL1* (rs1420101)^3,4,19^, and 3 non-synonymous coding SNPs located on the TIR domain (rs4988956, rs10192036 and rs10192157)^4^. Eight subjects carrying the low-risk and high-risk *IL-1RL1* haplotype were selected in each disease group. PBMCs isolated using Lymphoprep™ (Axis-Shield, Stemcell Technologies, #07801) and stored in liquid nitrogen were thawed, and resting CD4^+^ T cells were isolated with LS magnetic separator columns (Miltenyi Biotec, #130-096-533). Subsequently, CD4^+^ T cells were differentiated into Th2 cells using a human Th2 cell differentiation kit (CellXVivo, #CDK002) were harvested after 13 days.

### IL-33 stimulation of differentiated Th2 cells and RNA Sequencing

On day 13, Th2 cells were re-activated using 1μg/mL αCD3 (Purified NA/LE Mouse anti-Human CD3, 555329, BD Pharmingen) and 1μg/mL αCD28 (Purified NA/LE Mouse anti-Human CD28, 555725, BD Pharmingen) in the presence of either 0 or 100ng/mL IL-33 (rhIL-33, 3625-IL-010, BioTech). Total RNA was isolated after 24h of culture using TRIzol^®^Reagent (Life Technologies)^20^. RNA-seq libraries were prepared using NextFlex Directional qRNA-Seq kit (Bio Scientific). RNA sequencing was performed on an Illumina NextSeq500 platform (SBS50 kit, high-output mode, Illumina, San Diego, CA) using default parameters for paired-end sequencing (76bp + 9bp). Sequencing data was aligned to GRCh38 (Ensembl gene build 88) and quantified using STAR aligner, version 2.5.3a. Reads mapped to the same location and having identical UMIs were treated as PCR duplicates.

### Protein assays

Protein levels of IL-1RL1a, IL-4, IL-5, and IL-13 levels were determined in supernatants using Luminex at 72h after stimulation. Data were analysed using GraphPad Prism 8.3.1.

### Differential gene expression

After QC, differential gene expression (DGE) was analysed in R (Rstudio v3. 6.1) using Voom transformation in limma package^21^. Univariate (paired) analyses were performed to assess the effect of re-stimulation and IL-33 treatment, stratifying the samples based on *IL-1RL1* haplotype or disease status. The distribution of the fold changes in gene expression induced by IL-33 treatment was compared between groups using a T test. In all analyses, genes were considered statistically differentially expressed when the FDR adjusted p-value was <0.05. Gene ontology analysis was made using Shiny GO (0.76) software and verified using g-Profiler.

### Generation and analysis of gene signatures

Genes induced by T cell activation or IL-33 (FDR <0.05) were divided into 4 quartiles based on baseline gene expression levels. Gene signatures were generated using the top 5% genes with the highest LFC in the quartile of genes with the highest baseline gene expression. Gene set variation analysis (GSVA)^22^ was used to analyse enrichment scores (ES) for the gene signatures in U-BIOPRED sputum transcriptomics (GEO accession no. GSE76262)^23,24^. ES values were compared between patient groups with different severities^25^ or different sputum cell compositions^26^ and between U-BIOPRED Th2 based molecular phenotypes or transcriptome-associated clusters (TACs)^24^. Plots with Hommel^27^ multiple testing corrected Wilcoxon rank-sum test ^28^,statistical comparisons were generated using ggplot2^29^ and ggpubr R package^30^. Subjects were defined as High ES and Low ES based on the upper and lower tertiles of the ES values, compared for clinical parameters^25^ using Student’s T test ^31^ (normal distribution) or Wilcoxon rank-sum test^32^ for continuous data, and using Fisher’s exact test for categorical data^33^.

## Results

### Cohort characteristics

A total of 16 subjects carrying the asthma risk *IL-1RL1* haplotypes (based on rs1420101 (AA), rs4988956 (GG)), as well as 16 subjects carrying the protective *IL-1RL1* haplotype, were included in the study. For both haplotype groups, we used 8 asthma patients and 8 healthy subjects, resulting in an experimental design with 4 groups defined by a combination of disease state and *IL-1RL1* haplotype. PBMC donors were matched as much as possible, but the healthy control group contained fewer male participants compared to the asthma group, and healthy subjects were of younger age. General characteristics can be found in table 1.

**Table 1:** Study cohort characteristics.

| Features (Asthma) | High risk haplotype (8) | Low risk haplotype (8) |
| --- | --- | --- |
| Gender N (% male) | 3 (37.5%) | 7 (87.5%) |
| Age (years, mean) | 49.5 (2.2) | 50.2 (1.7) |
| FEV1 (L, prebronchodilator) mean (stdev) | 2.9 (0.5) | 3.2 (0.8) |
| FEV1/FVC (% , prebronchodilator) mean (stdev) | 69.6 (6.6) | 61.4 (12.6) |
| Blood eosinophils (10<sup>9</sup>/L) mean (stdev) | 0.21 (0.10) | 0.12 (0.08) |

| Features (Healthy) | High risk haplotype (8) | Low risk haplotype (8) |
| --- | --- | --- |
| Gender N (% male) | 1 (12.5%) | 3 (37.5%) |
| Age (years, mean) | 32.6 (14.5) | 36.8 (19.0) |
| FEV1 (L, prebronchodilator) mean (stdev) | 3.9 (0.9) | 3.6 (0.5) |
| FEV1/FVC (% , prebronchodilator) mean (stdev) | 82.8 (6.9) | 80.1 (2.9) |
| Blood eosinophils (10<sup>9</sup>/L) mean (stdev) | 0.21 (0.15) | 0.11 (0.06) |

FEV1 = the Forced Expiratory Volume in one second in litre (L) . FEV1/FVC = Forced Expiratory Volume in one second/Forced Vital Capacity ratio expressed in percentage. Mean (standard deviation. Medians (interquartile ranges) are shown unless otherwise stated.

### IL-33 induces T cell regulatory and activation pathways

Analysis of the transcriptomic changes induced by CD3/CD28 stimulation of Th2 cells using all subjects (n=29) revealed a total of 9299 genes (4687 up-regulated, 4312 down-regulated) that were significantly differentially expressed compared to unstimulated cells (figure 1a). Gene ontology (GO) analysis showed that genes in activated Th2 cells were involved in RNA processing, DNA replication and mitochondrial translocation elongation (figure 1b). No significant differences in CD3/CD28 induced genes were found between the *IL-1RL1* haplotype groups or the disease groups (supplemental figure 1).

**Figure 1.**
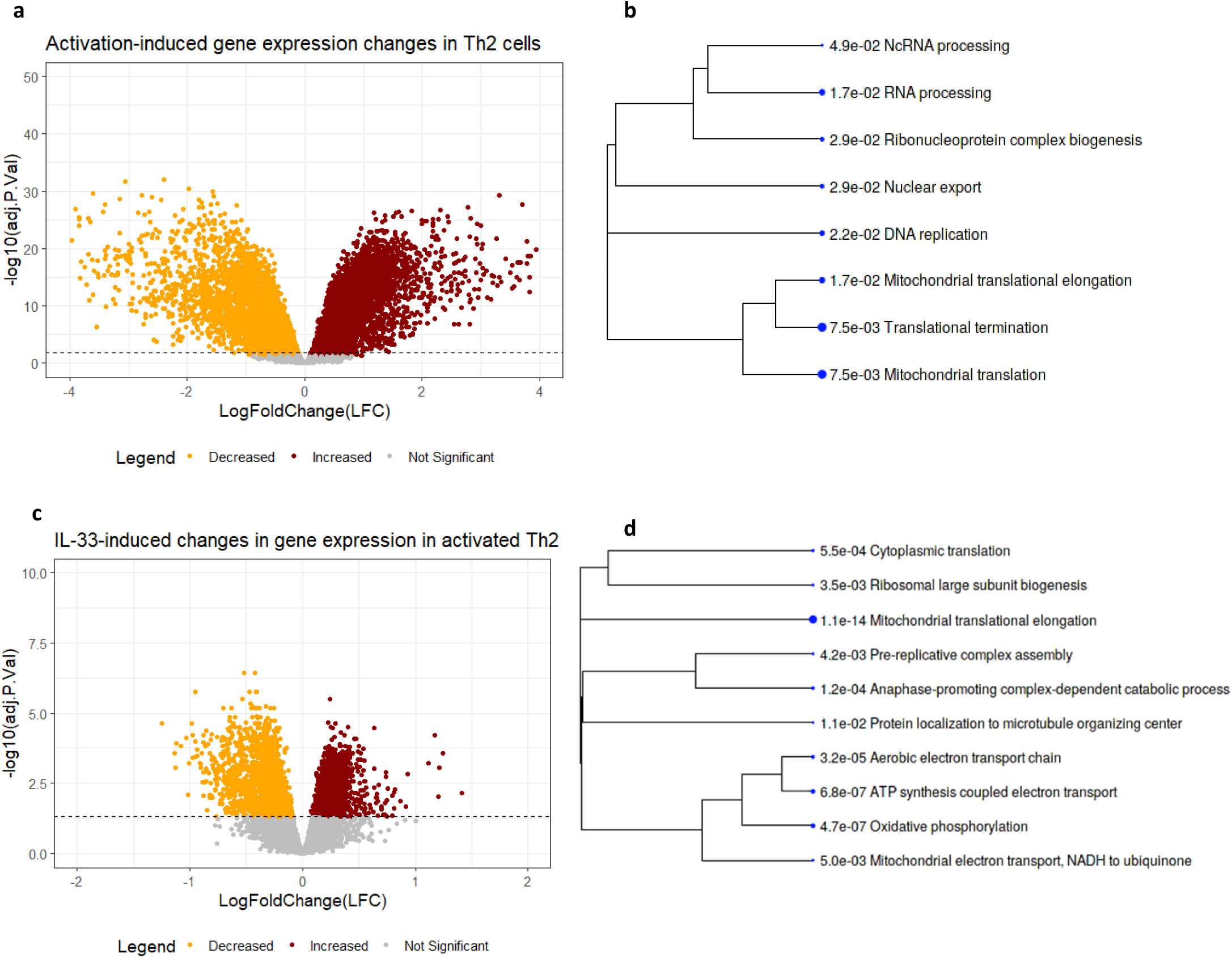
Differential gene expression of CD3/CD28 activated Th2 cells in the presence/absence of IL-33. ***a**) Volcano plot of FDR adjusted P-value against LogFoldChange (LFC) for the change in gene expression induced by Th2 cell activation in the total dataset (n=29)* with 12038 genes tested, significantly upregulated (brown dots) and downregulated (orange dots) genes are indicated. *Horizontal dotted black line represents FDR-adjusted p-value of 0.05. **b**) Hierarchical clustering (based on correlation among significant genes) of GO terms in genes enriched after Th2 cell activation (ShinyGO 0.76). Bigger dots indicate more significant FDR-corrected p values among the GO-pathways (biological pathways and molecular function). **c**) Volcano plot of gene expression changes induced by IL-33 stimulation of Th2 cells in the total dataset (n=29). **d**) Hierarchical clustering of GO terms enriched in IL-33-induced changes in gene expression*.

The presence of IL-33 induced prominent changes in the transcriptome of activated Th2 cells, with altered expression levels of 3677 genes (1568 up-regulated; 2078 down-regulated; figure 1c). GO analysis indicated that IL-33 treatment affected mitochondrial translation elongation and oxidative phosphorylation (figure 1d).

### IL-33 induced T cell activation is dependent on *IL-1RL1* haplotype and disease status

Next, we analysed the effect of *IL-1RL1* haplotype and disease status on the IL-33 driven transcriptional changes in activated Th2 cells. In all subjects (irrespective of disease status) carrying the risk haplotype of *IL-1RL1*, a large number of genes were differentially expressed between Th2 cells stimulated in the presence versus the absence of IL-33 (figure 2a, n=14). However, when analysing all subjects (irrespective of disease status) with the low-risk haplotype of *IL-1RL1*, IL-33 treatment did not induce any significant changes in gene expression (figure 2b, n=15). These data indicate that *IL-1RL1* haplotype is a strong determinant of the sensitivity of Th2 cells to IL-33.

**Figure 2.**
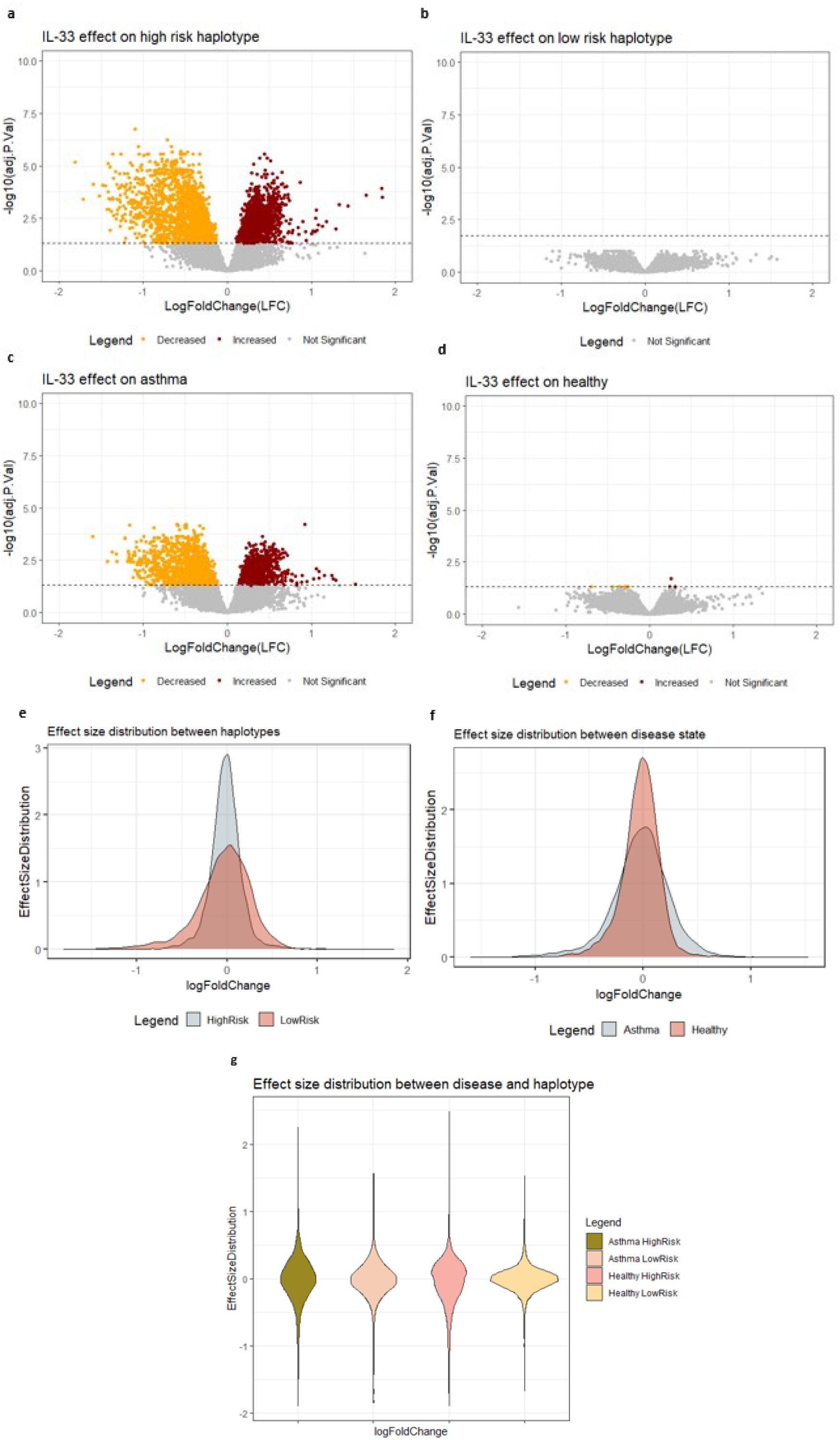
Effect of IL-33 treatment on gene expression of Th2 cells is driven by *IL-1RL1* haplotype and disease. *Volcano plots displaying the −log10 (FDR adjusted p-value) against log10 (LFC) for the changes in gene expression induced by IL-33 stimulation of Th2 cells, stratified on IL-1RL1 haplotype and disease status. Total number of genes tested*: 12038 *. Brown dots: upregulated genes. Orange dots: downregulated genes. **a**) high-risk haplotype carriers (n=14). **b)** low-risk haplotype carriers (n=15). **c)** asthma patients (n=15). **d)** healthy controls (n=14). Horizontal black dotted line represents the FDR-adjusted p-value =0.05. **e)** The distribution of Log10 (fold change) values for all genes induced by IL-33 stratified for haplotypes. (T-test; p-value = 1.09e-11). **f)** The distribution of Log10 (fold change) values for all genes induced by IL-33 stratified for disease groups. (T-test; p-value = 0.34). **g)** The distribution of all genes induced by IL-33 stratified for both disease and haplotype*.

In addition, we analysed the effect of disease status of the donors on the IL-33 induced responses in Th2 cells. In activated Th2 cells from patients with asthma, irrespective of their *IL-1RL1* haplotype, presence of IL-33 changed expression of 2524 genes (figure 2c). In contrast, only 10 genes were significantly changed by presence of IL-33 in Th2 cells isolated from healthy subjects, irrespective of their *IL-1RL1* haplotype (figure 2d). These data indicate that disease status also influences the sensitivity of Th2 cells for IL-33.

To directly test whether the effect of the *IL-1RL1* haplotype on IL-33 driven changes in gene expression of activated Th2 cells is different between patients with asthma and healthy controls, we analysed the interaction of haplotype and disease state on IL-33-induced changes in gene expression in activated Th2 cells (supplemental figure 2,3). We did not observe a significant interaction term, indicating that the effects of *IL-1RL1* haplotype are not different between the two disease groups and vice versa.

The lack of an interaction between disease state and IL1RL1 haplotype indicates that we mainly observe quantitative and not qualitative differences between the groups, and that magnitude of the response is determined by a combination of *IL-1RL1* haplotype and disease status. Therefore, to test whether the differences between the two haplotypes or the disease groups was reflected in the effect size of the response to IL-33, we next compared the distribution of the IL-33 induced fold changes in gene expression for all genes (including non-significantly differentially expressed genes) in the dataset between the two IL-1RL1 haplotypes (figure 2e) as well as between the two disease states (figure 2f). This analysis revealed a significantly larger IL-33 induced fold change of gene expression in *IL-1RL1* high-risk Th2 cells compared to low-risk carriers (T-test; p-value = 1.09e-11; figure 2e). In contrast, we did not observe a significant difference in the effect size of IL-33 induced gene expression in Th2 cells between the two disease groups (T-test; p-value = 0.34; figure 2f). Splitting the data into the 4 groups defined by the combination of disease status and IL-1RL1 haplotype, we observed that the healthy, low-risk group displayed a very narrow distribution of effect sizes in response to IL-33 compared to the other groups (figure 2g).

### *IL-1RL1* is differentially expressed upon IL-33 stimulation dependent of haplotype and disease status

Since we noticed an effect of both disease status and *IL-1RL1* haplotype on the Th2 response to IL-33, we hypothesized that the level of *IL-1RL1* expression differs between Th2 cells from the different groups. Analysing *IL-1RL1* expression levels in activated Th2 cells in the absence of IL-33 showed elevated levels of *IL-1RL1* expression in the low-risk haplotype (figure 3a), while *IL-1RL1* expression did not differ between disease groups. We see no differences between groups in the effect of IL-33 on *IL-1RL1* expression (figure 3b). Although, overall expression of *IL-1RL1* expression and numbers are induced by IL-33 irrespective of the disease state and haplotype (supplementary figure 4a). We also investigated protein expression of IL-33 stimulated Th2 cells in supernatants at 72 hours, including sST2, IL-4, IL-5 and IL-13, however, we did not observe an effect of IL-33 stimulation on IL-1RL1a expression or Th2 cytokine levels (supplemental figure 5).

**Figure 3.**
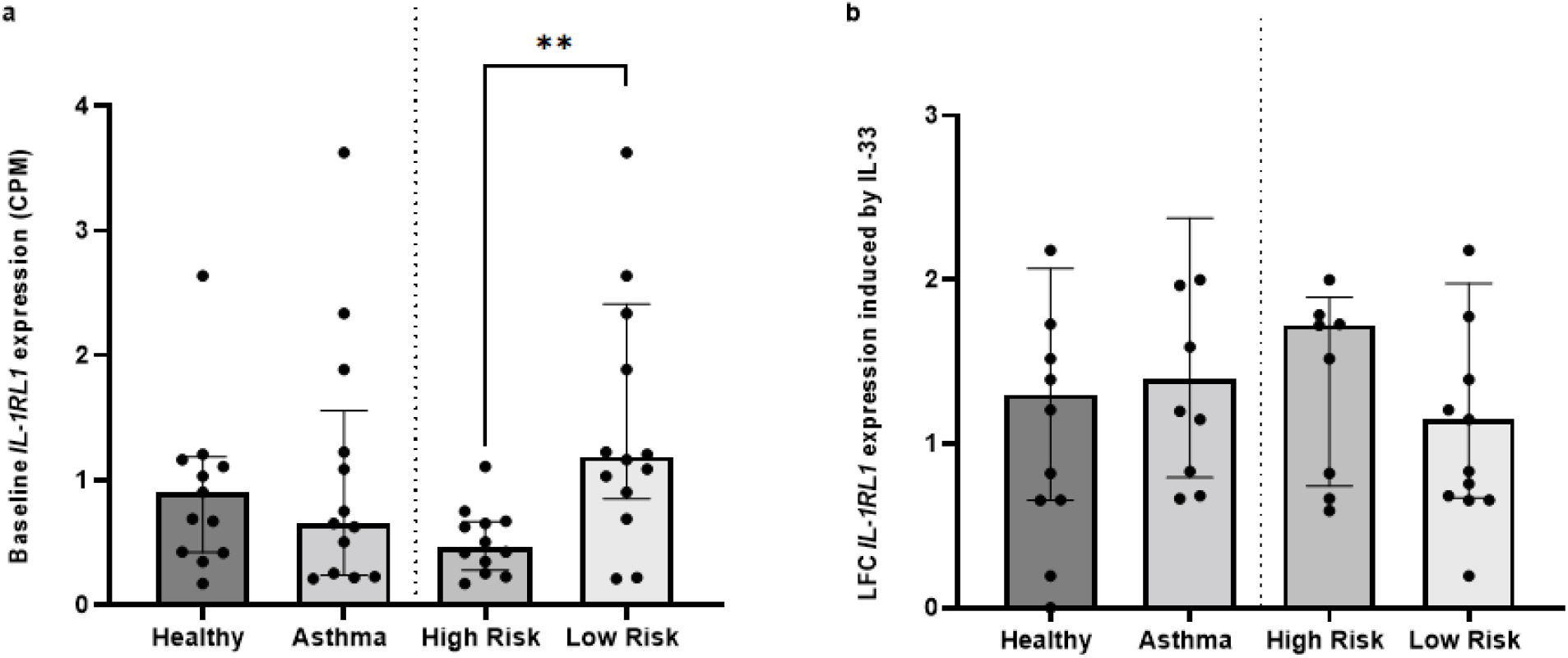
IL-33 induces *IL-1RL1* expression and numbers in Th2 cells to a larger extent in *IL-1RL1* risk haplotype carriers and asthma patients. **a)** normalized expression (counts-per-millions) of IL-1RL1 gene expression in activated Th2 cells in haplotype and disease groups (**, p=0.0025) **b)** log fold change of IL-1RL1 gene expression induced by IL-33 in activated Th2 cells in haplotype and disease groups.. Statistics: Panel a,b: Mann Whitney U test; *** p<0.001; ** p<0.01; *p<0.05.

### IL-33 induced Th2 gene signature shows strong correlation with pauci-granulocytic asthma

Given the critical role for Th2 cells in chronic inflammation in asthma, and the profound transcriptomic changes induced by IL-33 in activated Th2 cells, we hypothesized that a signature of activated Th2 cells or of IL-33 induced genes could be used in transcriptomic data from asthma patients to identify specific a subgroups of patients. To test this hypothesis, we generated gene signatures of IL-33 induced Th2 genes, by taking the top 5% of genes with the highest fold change within the subset of differentially expressed genes with baseline expression (in activated Th2 cells) in the highest quartile (supplementary figure 6a,b). Enrichment scores (ESs) calculated by GSVA for the IL-33 induced Th2 signatures when compared to other T-cell gene sets indicated a Th2-cell type specific response (supplementary figure 6c). IL-33-induced gene signatures (figure 4) were also calculated by GSVA in sputum transcriptomes from U BIOPRED study participants, and asthma patients with the highest 33% of ES scores were compared to those with the lowest ES scores. We found that patients with a high ES for the IL-33-induced gene signature have significantly better lung function, less severe asthma and lower eosinophil and higher macrophage counts in sputum (figure 4, supplementary figure 7). In addition, when comparing patient groups we find that the IL-33-induced gene signature showed a strong association with healthy controls, patients with pauci-granulocytic asthma and patients belonging to the TAC3 transcriptionally defined subgroup of asthma^24^ (figure 4). We replicated these analyses in the ADEPT replication cohort, and show that the enrichment score of Th2 and the IL-33 gene signatures show a similar trend for an association with the pauci-granulocytic phenotype of asthma (supplemental figure 8).

**Figure 4.**
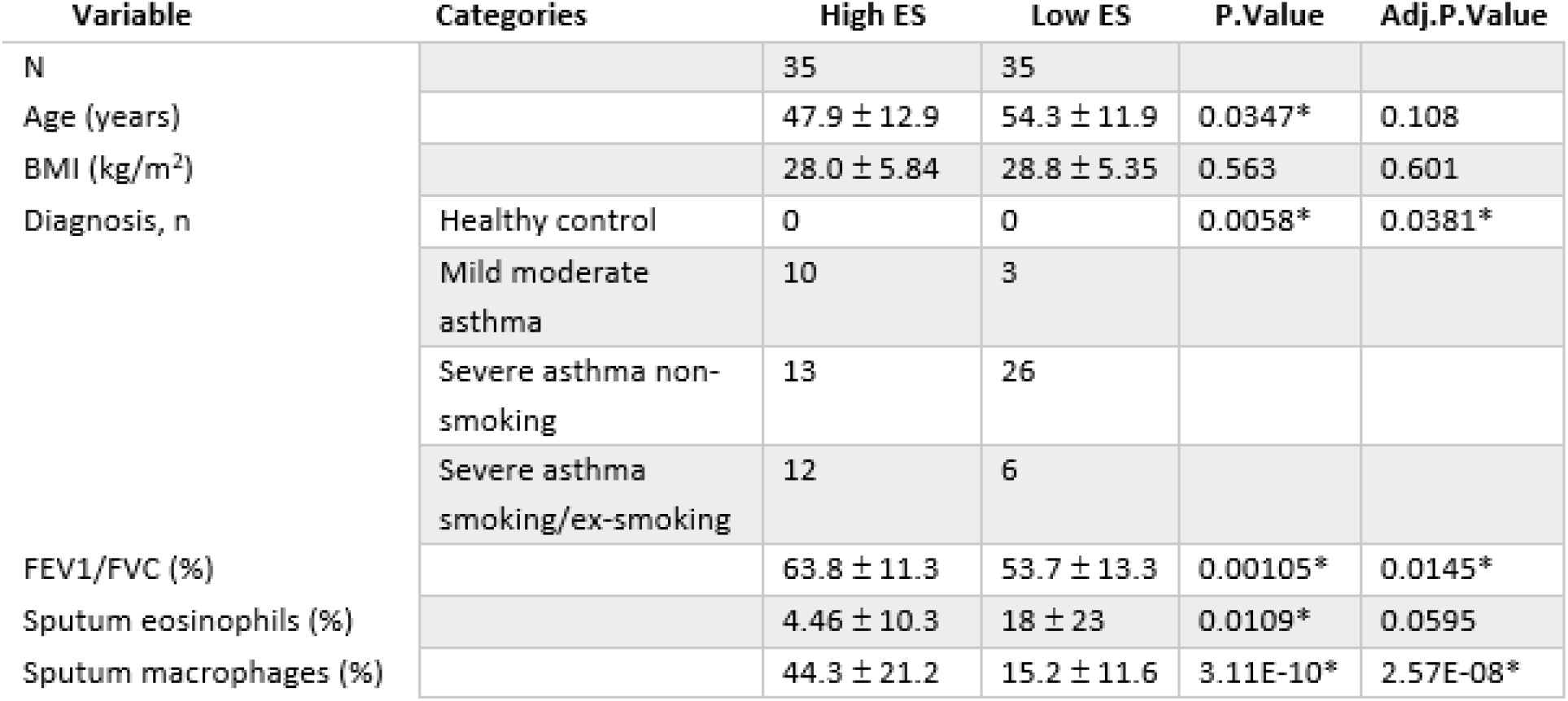
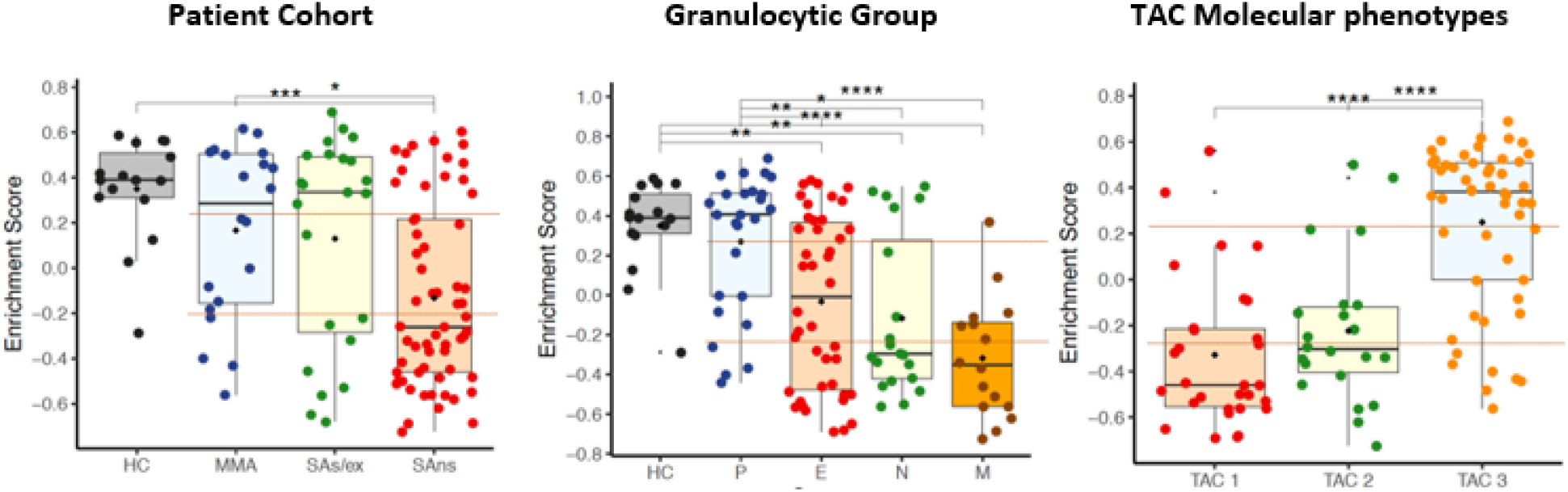
Gene signatures for Th2 activation and IL-33 activity show strong correlation with pauci-granulocytic asthma. **A)** Comparison of clinical variables between the upper versus the lower tertile of asthma patients in U BIOPRED based on the enrichment score for the IL-33 activation signature in sputum transcriptomic data (multiple testing corrected Wilcoxon rank-sum test)).B) Enrichment scores for the IL-33 gene signature in U BIOPRED sputum transcriptomic data plotted for the different disease groups (left), asthma phenotypes based on sputum cell counts (middle) and unsupervised clustering of U-BIOPRED patients based on their sputum transcriptomes (right, ^26^) using Student’s T test (normal distribution) or Wilcoxon rank-sum test for continuous data, and Fisher’s exact test for categorical data. *** p<0.001; ** p<0.01; *p<0.05

## Discussion

This study shows that IL-33 has a major impact on gene expression of activated Th2 cells, changing expression levels of around 25% of all genes detected. The IL-33 regulated genes are primarily involved in mitochondrial translational elongation and oxidative phosphorylation of the cells, which can be associated with mitochondrial metabolism in T cell differentiation^34^. In analysing the relevance of polymorphisms in *IL-1RL1* subunit of the IL-33 receptor complex to this response, we found that the *IL-1RL1* asthma risk haplotype resulted in a significantly larger magnitude of change in gene transcription induced by IL-33 compared to the protective allele. As a consequence, no differentially expressed genes were identified after IL-33 exposure in Th2 cells carrying the protective *IL-1RL1* allele. When contrasting disease states, while controlling for *IL-1RL1* haplotypes, we found that Th2 cells from asthma donors are also more sensitive to IL-33. We did, however, not observe an interaction between disease status of the donor and the *IL-1RL1* haplotype on the IL-33-induced response in activated Th2 cells, indicating that both IL1RL1 haplotype and disease status change the magnitude, but not the transcriptional profile, of the IL-33 induced response. *IL-1RL1* gene expression in activated Th2 cells was more strongly increased by IL-33 in asthma patients compared to healthy controls, while the *IL-1RL1* haplotype did not affect this. At the protein level, soluble IL-1RL1 levels in the supernatants of the Th2 cells, did not identify an effect of IL-33 stimulation, disease status or *IL-1RL1* haplotype. Finally, we generated a gene signature specific for IL-33 stimulated Th2 cells, that shows a strong association with the number of sputum macrophages in the U-BIOPRED severe asthma study and is relatively enriched in patients with pauci-granulocytic asthma and in patients belonging to the TAC3 cluster that was previously identified in U BIOPRED using unsupervised clustering of sputum transcriptomic data, and was characterised by metabolic and mitochondrial function genes^24^.

We are the first to show a significant effect of *IL-1RL1* asthma risk haplotype and disease status on IL-33 induced responses in primary Th2 effector cells. We defined the asthma risk haplotype based on two independent genetic signals: the rs1420101 SNP, a strong eQTL and pQTL for *IL-1RL1*, and three coding SNPs in complete linkage disequilibrium (LD): rs4988956, rs10192036 and rs10192157^3,4,19^. These latter 3 non-synonymous coding SNPs encode a functionally altered signalling domain in the cytoplasmic part of the IL-1RL1b receptor^4^. We did not observe higher expression of *IL-1RL1a* mRNA or protein in the Th2 cells from risk haplotype carriers, indicating that the enhanced signalling of IL-1RL1b intracellular domains might be the dominant effect regulating IL-33 responsiveness in Th2 cells. The effect of the *IL-1RL1* polymorphism and that of asthma status seems to be additive, since the asthma/risk *IL-1RL1* group showed the largest effect size of IL-33 on gene transcription, while asthma/protective *IL-1RL1* and healthy/risk *IL-1RL1* had a lower effect size, while in healthy/protective *IL-1RL1*, the effect size of IL-33 on gene transcription was the smallest. However, the IL-1RL1 haplotype effects seem to be dominant over the disease effects, as in the genotype-stratified analysis a significant difference was observed in IL-33 induced LFC, while in the reciprocal analysis, asthma donors did not show a significantly higher LFC in response to IL-33 (figure 2e,f). The effect of the asthma risk haplotype of IL-1RL1 might be explained by the presence of the risk TIR domain haplotype that we previously showed to be associated with increased signalling (using a NF-κB reporter) compared to the protective TIR domain haplotype in HEK293 cells ^35^).The mechanism of the disease effect on IL-33 induced signalling might be epigenetic regulation. We previously studied the potential link between asthma-associated SNPs at the IL-1RL1 locus, asthma disease status and *IL-1RL1* methylation patterns^19^. In this study, the T allele of the *IL-1RL1* SNP rs1420101 was found to be associated with lower serum IL-1RL1-a levels and higher blood eosinophil counts. Interestingly, variation in the protein levels were associated with blood DNA-methylation at different CpG sites in the *IL-1RL1* gene, and was associated with more pronounced allergic inflammation and higher asthma risk^36^. These data suggest that epigenetic programming in asthmatic donors could also have an effect on the Th2 cells upon activation^19^. However, this study was performed in whole blood, while we studied *ex vivo* cultured Th2 cells which are only a minor fraction of blood cells..

Given the importance of the IL-33/IL-1RL1 pathway in asthma and the ongoing clinical development of antagonists acting on this pathway^13,37,38^, we hypothesized that we could use a signature of IL-33-induced genes in Th2 cells to identify patients with relatively higher activity of this pathway based on RNA-sequencing data. Given that Th2 cells are a relatively rare cell population in airway wall, we selected the sputum transcriptome data of the U-BIOPRED severe asthma study to test our hypothesis^24^. We observed that the IL-33-induced Th2 gene signature was not enriched in sputum transcriptomic data from asthmatic patients when compared to healthy controls. Within asthma patients, our signature was associated with pauci-granulocytic asthma, with the previously identified TAC3 phenotype, and was highly correlated to the number of sputum macrophages. This was somewhat unexpected, as the IL-33/IL-1RL1 pathway has been repeatedly linked to type 2 high, eosinophilic asthma^24^. The strong positive correlation of the IL-33-induced Th2 signature with the fraction of sputum macrophages, however, may indicate an overlap in the transcriptional profile of Th2 cells and sputum macrophages. This might explain the association with pauci-granulocytic asthma and with the TAC3 subset of patients, which was reported to have relatively high numbers of sputum macrophages within the U-BIOPRED cohort^39^. As a role for sputum macrophages in asthma pathogenesis is currently unclear^40^, it remains to be determined whether this signature can be of use to identify a true asthma endotype. However, current data suggest that M1 macrophages are associated with disease progression and airway remodelling, while, M2 macrophages are associated with type-2 asthma^41^. Further work would need to test whether our signature is specific for either macrophage subset.

We need to also consider the strengths and weaknesses of our approach to test the functionality of asthma-associated genetic variation at the IL-1RL1 locus by using primary immune cells from asthma patients and controls. One strength of our approach is, that we perform our experiments using endogenous expression levels of the IL-1RL1 receptor, allowing us to focus on physiologically relevant effects on gene transcription induced by IL-33. Moreover, the naturally occurring haplotypes reflect true combinations of genetic variation and therefore incorporate potential SNP-context effects as compared to SNP mutation strategies that do not. It is well-known that the LD pattern at the *IL-1RL1* locus is complex^3,19^, and the SNPs that we have selected in the *IL-1RL1* locus may in fact be also affect expression levels of neighbouring genes (e.g. *IL1R1, IL18R1, IL18RAP*) with which these SNPs are also in high LD. Therefore, studying these SNPs in populations with a different LD pattern compared to the Caucasian-origin donors we have used, as was done recently in the African American population, might also provide an important way forward^42^. Moreover, we realize that, although we carefully tried to match the disease and haplotype groups on clinical parameters, there were fewer males in the healthy group, which may act as a potential confounder. Unfortunately, considering our stratified analyses we had limited numbers of donors to further study sex effects separately. Future studies would benefit from investigating isoform specific effects of *IL-1RL1* SNPs on *IL-1RL1* expression, as RNA differences in the variant encoding the soluble decoy IL-1RL1a versus the transmembrane form IL-1RL1b may have different mechanistic implications.

In conclusion, the current study shows that IL-33 strongly changes the transcriptional phenotype of activated Th2 cells in subjects carrying asthma-associated risk genotypes in *IL-1RL1* and in asthma patients. An IL-33-induced Th2 activated gene expression signature could identify a subset of patients with pauci-granulocytic phenotype of asthma and their corresponding TAC3 group of patients. This indicates that gene signatures need to be generated using cells present in sufficiently high numbers in the sample of interest which would benefit proper stratification of patients for anti-IL-33 targeted therapies.

## Acknowledgements

This study was supported by the Lung Foundation Netherlands (grant no. 3.2.09.081JU) and an investigator-initiated grant by GSK. We would like to thank M. Nieuwenhuis for her support in patient recruitment, as well as C. Vermeulen, B. Ditz, A. Faiz and M. Berg for their feedback during development of the analysis scripts.

## Abbreviations

ADEPT: Airway Disease Endotyping for Personalized Therapeutics
DEGs: Differentially expressed genes
ES: Enrichment score
E: Eosinophilic
FC: Fold-change
FEV1: Forced Expiratory Volume in the first second
FDR: False discovery rate
FVC: Forced Vital Capacity
GSVA: Gene Set Variation Analysis
HC: Healthy control
IL: Interleukin
IL-1RL1: Interleukin 1 Receptor Like 1
IL-33: Interleukin – 33
LFC: Log Fold Change
MMA: Mild-moderate asthma
M: Mixed
N: Neutrophilic
PBMC: Peripheral Blood Mononuclear Cells
P: Pauci-granulocytic
SA: Severe asthma
SAs/ex: Severe asthmatic smoker/ex-smoker
SNP: Single nucleotide polymorphisms
T2: Type 2
TAC: Transcriptome-Associated Cluster
Th2: T helper 2 lymphocyte
U-BIOPRED: Unbiased Biomarkers for the Prediction of Respiratory Disease Outcomes

## Notes

### Competing Interest Statement

The following authors have no conflict of interest to disclose: LH, YEB, NZK, SB, FND, GV, JV; IA declares grant support from EU-IMI, GSK, MRC, EPSRC; consulting fees from GSK, Sanofi, Chiesi, Kinaset and lecture fees from AZ, Sanofi, Eurodrug and Sunovion and payment for expert testimony Chiesi and support for travel from AZ, MvdB declares grant support from Chiesi, AZ, Novartis, Genentech and Roche, GHK declares grant support from Lung foundation Netherlands, TEVA the Netherlands, EU H2020 program, Ubbo Emmius Foundation, Vertex, consulting fees from AZ, Pure IMS, and lecture fees from Sanofi, Genzyme, MCN declares grant support from GSK, Lung Foundation Netherlands and the Netherlands Ministry of Economic Affairs and Climate Policy by means of the PPP-allowance; GHK, AKSJ, LH, MvdB, MEK declare grant support from GSK, Lung Foundation Netherlands. COI forms for ND, IS, SB, and YEB will be provided.

