## Supplemental Figure 1 for "IL-33 induced gene expression in activated Th2 effector cells is dependent on IL-1RL1 haplotype and disease status"

**Figure Information for:**

**Figure 1**

**a**

**b**

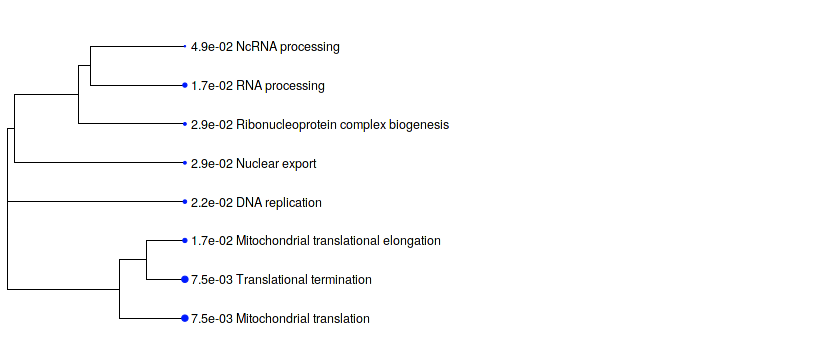

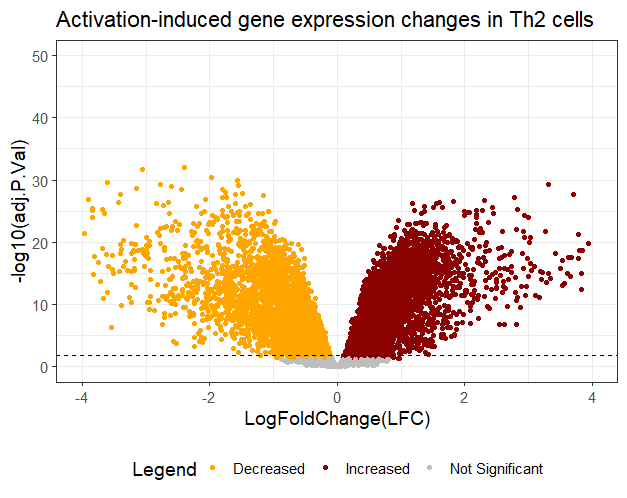

**c**

**d**

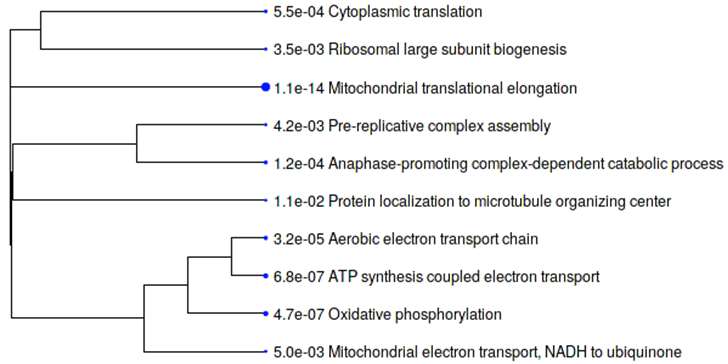

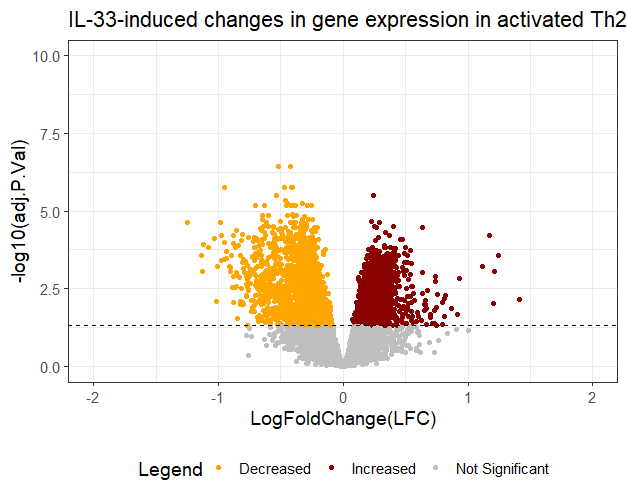

**Differential gene expression of CD3/CD28 activated Th2 cells in the presence/absence of IL-33. *a****)* *Volcano plot of FDR adjusted P-value against LogFoldChange (LFC) for the change in gene expression induced by Th2 cell activation in the total dataset (n=29)* with 12038 genes tested, significantly upregulated (brown dots) and downregulated (orange dots) genes are indicated. *Horizontal dotted black line represents FDR-adjusted p-value of 0.05.* ***b****) Hierarchical clustering (based on correlation among significant genes) of GO terms in genes enriched after Th2 cell activation (ShinyGO 0.76). Bigger dots indicate more significant FDR-corrected p values among the GO-pathways (biological pathways and molecular function).* ***c****) Volcano plot of gene expression changes induced by IL-33 stimulation of Th2 cells in the total dataset (n=29).* ***d****) Hierarchical clustering of GO terms enriched in IL-33-induced changes in gene expression.*
