## Supplemental Table 1 for "IL-33 induced gene expression in activated Th2 effector cells is dependent on IL-1RL1 haplotype and disease status"

**Table Information for:**

**Table 1: Study cohort characteristics**

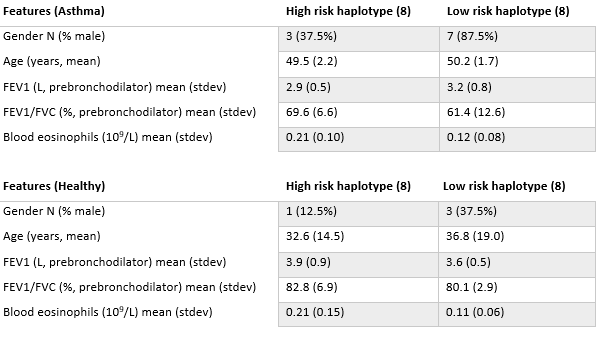

FEV1 = the Forced Expiratory Volume in one second in litre (L) . FEV1/FVC = Forced Expiratory Volume in one second/Forced Vital Capacity ratio expressed in percentage. Mean (standard deviation. Medians (interquartile ranges) are shown unless otherwise stated.
