## Supplemental Figure 2 for "IL-33 induced gene expression in activated Th2 effector cells is dependent on IL-1RL1 haplotype and disease status"

**Figure Information for:**

**Figure 2**

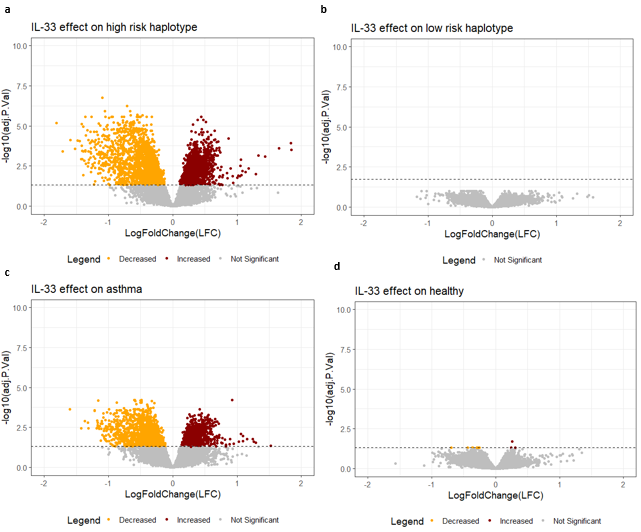

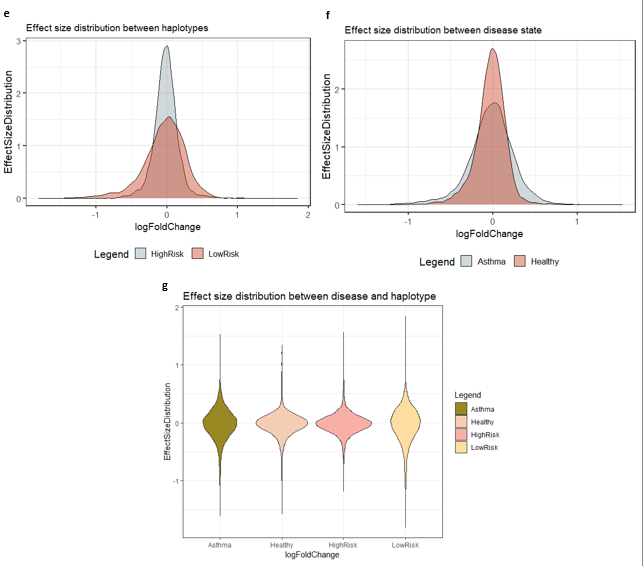

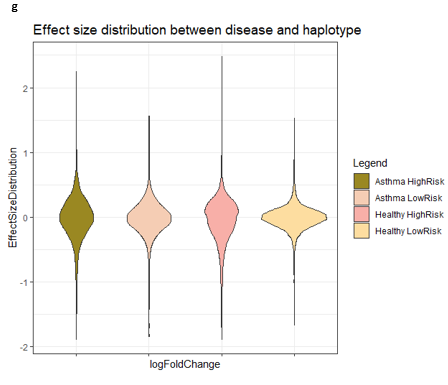

**Effect of IL-33 treatment on gene expression of Th2 cells is driven by *IL-1RL1* haplotype and disease.** *Volcano plots displaying the -log10 (FDR adjusted p-value) against log10 (LFC) for the changes in gene expression induced by IL-33 stimulation of Th2 cells, stratified on IL-1RL1 haplotype and disease status. Total number of genes tested:* 12038 *. Brown dots: upregulated genes. Orange dots: downregulated genes.* ***a****) high-risk haplotype carriers (n=14).* ***b)*** *low-risk haplotype carriers (n=15).* ***c)*** *asthma patients (n=15).* ***d)*** *healthy controls (n=14). Horizontal black dotted line represents the FDR-adjusted p-value =0.05.* ***e)*** *The distribution of Log10 (fold change) values for all genes induced by IL-33 stratified for haplotypes. (T-test; p-value = 1.09e-11).* ***f)*** *The distribution of Log10 (fold change) values for all genes induced by IL-33 stratified for disease groups. (T-test; p-value = 0.34).* ***g)*** *The distribution of all genes induced by IL-33 stratified for both disease and haplotype.*
