## Supplemental Figure 3 for "IL-33 induced gene expression in activated Th2 effector cells is dependent on IL-1RL1 haplotype and disease status"

**Figure Information for:**

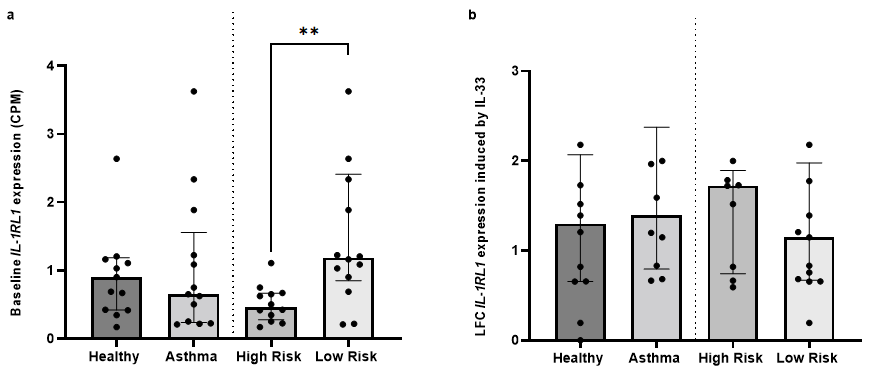
**Figure 3**

**I****L-33 induces *IL-1RL1* expression and numbers in Th2 cells to a larger extent in *IL-1RL1* risk haplotype carriers and asthma patients. *a)*** *normalized expression (counts-per-millions) of IL-1RL1 gene expression in activated Th2 cells in haplotype and disease groups (**, p=0.0025)* ***b)*** *log fold change of IL-1RL1 gene expression induced by IL-33 in activated Th2 cells in haplotype and disease groups.. Statistics: Panel a,b: Mann Whitney U test; *** p<0.001; ** p<0.01; *p<0.05.*
