## Supplemental Figure 4 for "IL-33 induced gene expression in activated Th2 effector cells is dependent on IL-1RL1 haplotype and disease status"

**Figure Information for:**

**Figure 4**

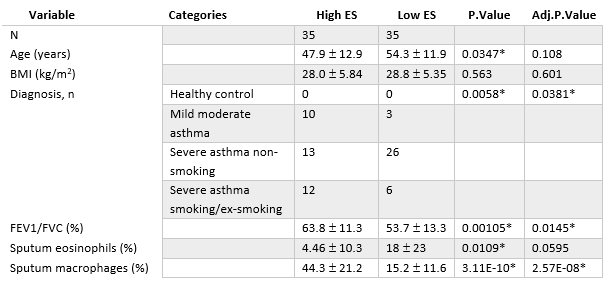

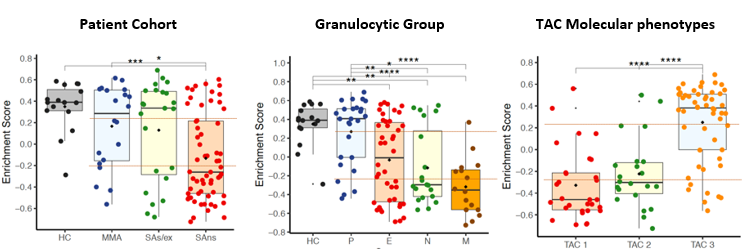

**Gene signatures for Th2 activation and IL-33 activity show strong correlation with pauci-granulocytic asthma. *A)*** *Comparison of clinical variables between the upper versus the lower tertile of asthma patients in U BIOPRED based on the enrichment score for the IL-33 activation signature in sputum transcriptomic data (multiple testing corrected Wilcoxon rank-sum test) ).B) Enrichment scores for the IL-33 gene signature in U BIOPRED sputum transcriptomic data plotted for the different disease groups (left), asthma phenotypes based on sputum cell counts (middle) and unsupervised clustering of U-BIOPRED patients based on their sputum transcriptomes (right,* ^26^*) using Student’s T test (normal distribution) or Wilcoxon rank-sum test for continuous data, and Fisher's exact test for categorical data .* **** p<0.001; ** p<0.01; *p<0.05*
